# Clinically Relevant Biomarkers of Alzheimer’s Disease Are Associated with Age and Cortical Atrophy in Chimpanzees (*Pan troglodytes*)

**DOI:** 10.64898/2026.08.12.744516

**Authors:** Michele M. Mulholland, Elizabeth R. Magden, Angela M. Achorn, Jean-Francois Mangin, William D. Hopkins

## Abstract

Chimpanzees share a number of age-related brain changes with humans, such as reductions in neurons and increases in neuropathology. To date, there are no published studies of peripheral biomarkers related to Alzheimer’s pathology and their associations with age and cortical atrophy in chimpanzees. Here we examined cross-sectional differences and longitudinal changes in biomarkers of pathological protein aggregation, neuroinflammation, and microglial function measured in serum. We examined the relationships between biomarkers and clinically relevant biomarker ratios with both age and cortical atrophy. We found linear and quadratic relationships between age and several biomarkers and ratios. Most biomarkers increased with age. While controlling for sex, we found significant negative associations between age and sulci surface area, mean depth, and gray matter thickness and a positive association with fold opening. Aβ42 and Aβ40 showed higher biomarker values associated with lower surface area, mean depth, and gray matter thickness and higher fold opening values. The clinically relevant biomarker ratios were also associated with cortical atrophy – Aβ42/Aβ40 was negatively associated with gray matter thickness, and pTau217/Aβ42 (both total and brain-derived) was positively associated with surface area and gray matter thickness and negatively associated with fold opening. Consistent with our hypotheses and previous findings in humans, many peripheral biomarkers associated with neurodegeneration and Alzheimer’s disease increase as chimpanzees age. We believe this is the first evidence demonstrating an association between these clinically relevant biomarkers of Alzheimer’s disease and phenotypes of brain aging in nonhuman primates, underscoring their importance as models of aging and neurodegenerative disease.

## Introduction

Genetically and phylogenetically, chimpanzees (*Pan troglodytes*) are one of two extant species most closely related to humans. Additionally, like other nonhuman primates, chimpanzees share a number of physiological, neurological, behavioral and cognitive abilities with humans that are absent or less developed in other species that serve as models in behavioral and biomedical research ^1–4^. A specific case in point is research on the comparative biology of aging, including Alzheimer’s disease (AD) and related dementias ^5,6^. Though shorter-lived than humans, chimpanzees and other great apes have the longest healthspans and lifespans among nonhuman primates ^7^ with some evidence suggesting that females live, on average, 7 years longer than males ^8–10^. Furthermore, like humans, previous studies have reported that chimpanzees show increasing atrophy of the brain and age-related decline in whole brain and region-specific gray matter volume and cortical thickness ^11–16^. Chimpanzees also show cross-sectional and longitudinal changes in cognitive functions as well as slowing in motor functions including tool use proficiency ^13,17–22^.

Regarding neuropathology and aging, geriatric chimpanzees (and other great apes) co-express both amyloid beta plaques (Aβ) and neurofibrillary tangles, two common features found in postmortem brains of patients with probable AD ^23–27^. Additionally, like in humans, cerebral amyloid angiopathy is frequently observed in geriatric chimpanzee brains ^23^. There is also some evidence that aged chimpanzees show loss in neurons within regions of the hippocampus, and particularly among individuals that exhibit some form of neuropathology ^28^. Thus, data on brain aging in chimpanzees ^29–31^ suggest that there are many similarities to humans.

The purpose of this study was two-fold. First, we tested for longitudinal changes and cross-sectional associations between age and serum biomarkers of neurodegeneration in a sample of chimpanzees. Studies in humans have shown that fluid biomarkers of AD and related dementias such as Aβ40, Aβ42, total tau and phosphorylated tau can be detected from CSF, plasma, and serum and these measures correlate with loss in cognitive functions ^32–36^ and AD neuropathology using either PET imaging ^37–39^ or direct measures obtained from postmortem brains ^40–43^. Associations between age and biomarkers of neurodegeneration have previously been reported in several nonhuman primate species including marmosets, squirrel monkeys, vervet monkeys, rhesus monkeys, cynomolgus monkeys and baboons ^5,8,44–47^ but we know of none in chimpanzees.

We used the multiplex NULISAseq CNS Disease Panel 120 which measures >120 biomarkers broadly classified into eight categories of CNS function including (1) pathological protein aggregation, (2) synaptic and neuronal network defects, (3) aberrant proteostasis, (4) cytoskeletal abnormalities, (5) altered energy homeostasis, (6) DNA and RNA defects, (7) inflammation, and (8) neuronal cell death. In this study, we were specifically interested in the subset of biomarkers within the protein pathological aggregation category which included amyloid beta (Aβ38, Aβ40, Aβ42), phosphorylated tau (pTau181, pTau217 and pTau231 and their associated brain-derived counterparts: BD-pTau181, BD-pTau217, BD-pTau231), Huntingtin (HTT), total tau (MAPT and BD-MAPT), beta-site APP-cleaving enzyme (BACE1), presenilin-1 (PSEN1), alpha-synuclein phosphorylated at serine-129 (pSNCA-129), alpha-synuclein (SNCA), beta-synuclein (SNCB), and the phosphorylated form of the TAR DNA binding protein 43 (pTARDBP-S409).

We also characterized three additional markers previously examined in human and nonhuman primates that are associated with neuroinflammation and microglial function including neurofilament light (NfL), glial fibrillary acidic protein (GFAP), and triggering receptor expressed on myeloid cells 2 (TREM2) ^42,44^. In addition to the individual biomarkers, two biomarker ratio measures have been shown to distinguish between patients expressing different dementia severity or at different clinical stages of AD, Aβ42/Aβ40 and pTau-217/Aβ42 ^34,35^. Here we examined these two ratios along with brain-derived pTau17/Aβ42. Based on the scaling of these ratio measures and their association with age and disease status in humans, we hypothesized that older chimpanzees would have lower Aβ42/Aβ40 and higher pTau-217/Aβ42 values (including for the brain-derived pTau217). Based on the existing neuropathology findings in chimpanzees ^23–25^ and other nonhuman primates, we hypothesized that age would significantly and positively correlate with one or more of the biomarkers. For the cross-sectional analyses, we tested for both linear and quadratic associations between age and the biomarkers.

In a second set of analyses, we examined whether the biomarkers of neurodegeneration were associated with measures of cortical atrophy in the chimpanzees. In humans, the brain becomes increasingly atrophied in older individuals as evident by declines in brain weight, gray and white matter volume, surface area, cortical thickness, and increasing size of the ventricles ^48–50^. The morphological changes in the brain associated with age are hypothesized to be related to decline in microstructural organization such as loss of neurons, synaptic density, and demyelination of the cortex. In neurodegenerative disorders, such as AD or fronto-temporal dementia, brain atrophy can be accelerated and particularly severe due to neuropathological changes such as the accumulation of amyloid plaques and/or neurofibrillary tangles in cortex ^51–53^. As noted above, chimpanzees show age-related changes in various aspects of cortical morphology and here we tested for their association with the fluid serum biomarkers after accounting for sex and age. Of specific interest were associations between the biomarkers and measures of variation in 23 primary sulci including their surface area, average depth, gray matter thickness, and fold opening. Previous studies in humans, chimpanzees, and baboons have reported that increasing age is associated with declines in gray matter volume, surface area, gray matter thickness, and increases in fold opening ^15,54–56^; if biomarker values underlie individual variation in cortical atrophy, we hypothesized that the brain outcome measures would be associated with one or more biomarker measures after controlling for sex and age.

## Methods

### Subjects

Previously banked cross-sectional serum samples were analyzed from a total of 234 chimpanzees including 94 males and 140 females ranging age between 8.75 and 58.79 years (*Mean* = 27.82 years, *SD* = 11.21). Archived magnetic resonance image (MRI) scans were available from a subset of 225 chimpanzees including 89 males and 136 females. Serum samples and MRI scans were obtained prior to the 2015 implementation of changes in NIH policy on the use of chimpanzees in research. MRI scans and blood samples (up to 6 ml) were collected from the chimpanzees during their annual physical exams. Multiple aliquots of serum were derived from the blood samples and subsequently archived and stored in a −80 deg C freezer. In this report, we selected the archived serum sample that was collected at the same or closest time to the date of the chimpanzee’s MRI scan. The T1-weighted MRI scans were collected on either a Siemens 3T or 1.5T GE scanner and the methods and scanning sequences have been described in previous publications ^15,57^. Finally, in a subset of 24 chimpanzees (14 females, 10 males), we analyzed a second set of serum samples at an older age as a means of assessing longitudinal changes in the biomarkers of interest. These 24 chimpanzees ranged in age between 27 and 43 years at the time of the initial serum sample collection. The second sample was collected between 2 and 11 years after the initial sample (*Mean* = 6.65 years, *SD* = 3.02). At the time of serum sample collection and MRI scanning, all chimpanzees were housed at either the Emory National Primate Research Center (n=86) or the University of Texas MD Anderson Cancer Center (n=148).

### Cortical Atrophy

Archived T1-weighted MRI scans were resampled at .645 mm^3^, aligned in the ACPC plane, skull-stripped, bias correlated, and denoised following methods described in Hopkins, et al. ^58^. The scans were then imported into the software program BrainVisa for analysis. The scans were processed through the Morphologist pipeline and the sulci were extracted from the cortex using methods that have been described in detail for both human and nonhuman primate MRI scans ^56,59^. Shown in Figure 1A are 3D renderings of a chimpanzee brain showing the sulci extraction steps and cortical sulci labelled in this study ^60,61^. From the 3D renderings, 23 sulci were manually labeled including (1) central, (2) superior precentral, (3) inferior precentral, (4) superior frontal, (5) middle frontal, (6) inferior frontal, (7) fronto-orbital, (8) orbital, (9) lunate, (10) occipital temporal lateral, (11) occipital lateral, (12) occipital temporal medial, (13) inferior temporal, (14) middle temporal, (15) superior temporal, (16) sylvian fissure, (17) intraparietal, (18) superior postcentral, (19) inferior postcentral, (20) medial parietal occipital, (21) superior parietal, (22) cingulate, and (23) calcarine. For each sulcus, measurements of surface area (SA, mm^2^), mean depth (MD, in mm), gray matter thickness averaged across both sides (GMT, in mm), and fold opening (FO, in mm) were obtained for every subject in their native space (see Figure 1B). Because the MRI scans were acquired on scanners with different magnetic fields, we converted the SA, MD, GMT, and FO values for each sulcus to standardized *z*-scores within the cohorts of chimpanzees scanned on the 1.5T and 3T scanners. To limit the number of statistical analyses and guard against Type I error, rather than analyze each sulcus, we averaged the *z*-scores across the 23 sulci for SA, MD, GMT and FO measures for each subject. The mean *z*-scores were the primary outcome measures of interest.

**Figure 1.**
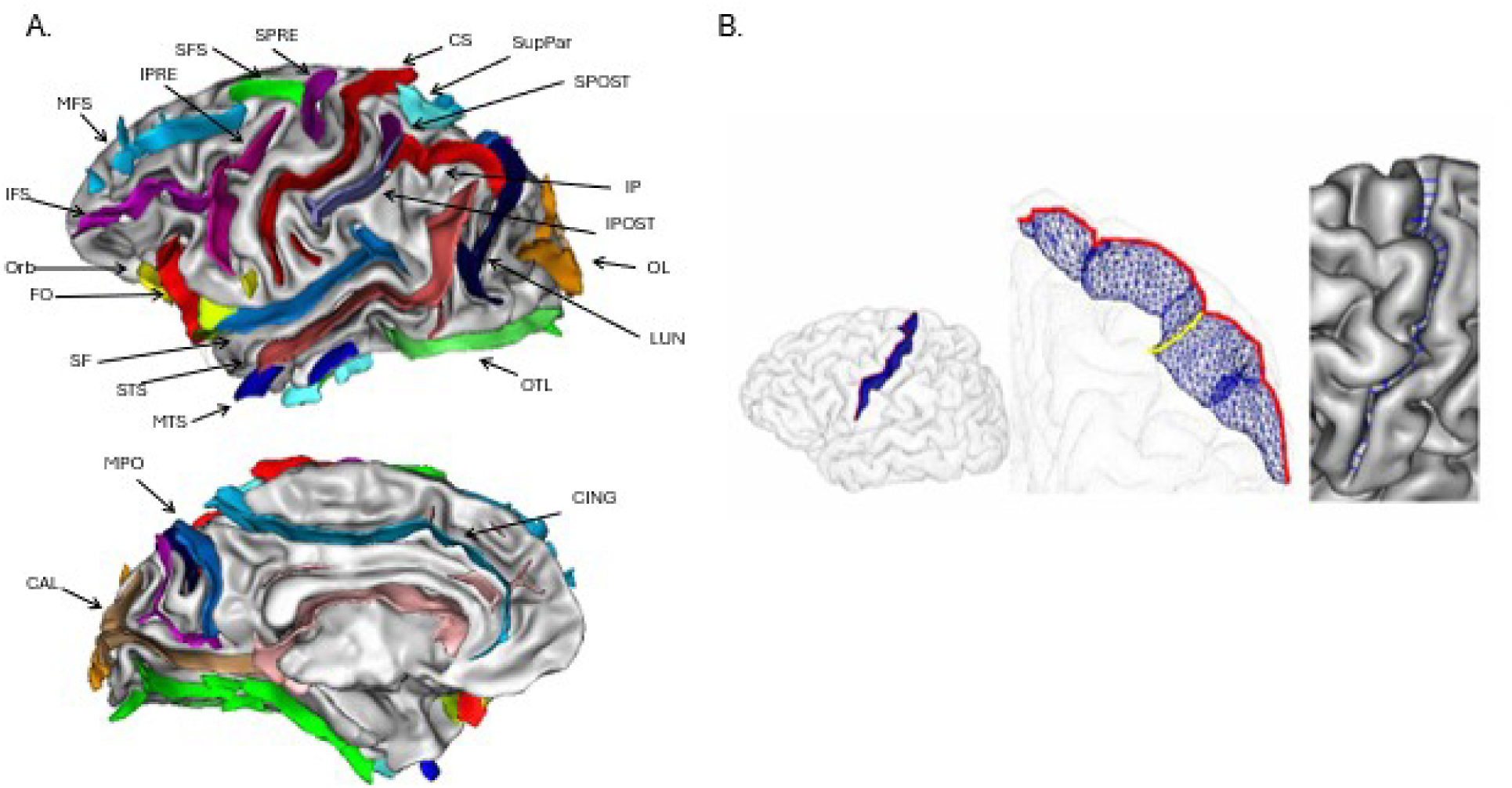
(A.) 3D renderings of a chimpanzee brain showing the sulci extraction and labeling in BrainVisa. Top: CS = central sulcus, IPRE = inferior precentral, SPRE = superior precentral, SFS = superior frontal, MFS = middle frontal, IFS= inferior frontal, Orb = orbital, FO = fronto-orbital, SF = sylvian fissure, STS = superior temporal, MTS = middle temporal, ITS = inferior temporal, IPOST = inferior post-central, SPOST = superior post-central, IP = intraparietal, SupPar = superior parietal, LUN = lunate, OL = occipital lateral, OTL = occipital temporal lateral. Bottom: MPO = medial parietal occipital, CING = cingulate, CAL = calcarine, OTM = occipital temporal medial. (B.) Diagram showing the SA (red), MD, GMT, FO (yellow) measurements that were obtained for each sulcus from every subject.

### Biomarker Analysis

Serum samples were analyzed using the NULISAseq CNS 120 disease panel (Next-generation Ultra-sensitive Ligand-based Immunoassay; Alamar Biosciences) proteomics platform through the MD Anderson-supported Oncology Research and Immuno-mONitoring (ORION) Core. Specifically, the NULISA platform uses immunoassay technologies with ultra-high sensitivity and dynamic range for protein biomarkers extracted from biological fluids. NULISAseq is a multiplex assay that employs quantification of proteins relative to a chosen standard and the values are expressed as units of fold change rather than absolute values (i.e., pg /mL). The primary outcome measures from the NULISA platform are NULISA Protein Quantification (NPQ), an Alamar Biosciences-specific relative quantification unit for this assay. NPQ is calculated by normalizing NGS read counts by an internal control, normalizing using interplate controls, rescaling the data, and finally, performing a log2 transformation. A one unit increase in NPQ represents a doubling of protein target level.

### Data Analysis

We know of no previously published data on neurodegeneration biomarkers in chimpanzees using the NULISAseq CNS 120 disease panel, so we have provided the quality control cross-reactivity percentages for the chimpanzees to the human markers across all proteins in Supplemental Table 1 as well as their associations with age for the entire sample. In Supplemental table 1, we distinguish between markers that were detectable, detectable but at levels below the minimum threshold established for the human standard, and undetectable (i.e., the majority of NPQ values were zero). There were only two biomarkers (ICAM1 and SAA1) which were not detectable. Though a majority of the subjects had NPQ values, an additional twelve biomarkers were deemed below threshold based on human standards according to quality control analyses of NULISA. The remaining biomarkers were all reliably detected, including > 95 % for the main biomarkers of interest in this study. For each biomarker of interest, the NPQ output data were initially analyzed for outliers using boxplots and individual data points outside the 99^th^ percentile, which were excluded from all subsequent analyses. Based on these criteria, outliers were very infrequent for the biomarkers of interest (fewer than 8 individuals for any given marker). The remaining NPQ values were analyzed using parametric statistics including analysis of covariance, regression, and partial correlation coefficients. Because our hypotheses were unidirectional, alpha was set to *p* < .05. In addition to the raw NPQ values, we computed three biomarker ratio measures that are frequently used in studies with humans and have been reported to be associated with age and identify subjects at risk for the development of AD ^34,35^. The three ratios were Aβ42/Aβ40, pTau217/Aβ42 and BD-pTau217/Aβ42. Because the NPQ values were already derived from logged raw biomarker values, we could not compute a true ratio but, instead, subtracted the numerator from denominator terms for each measure. Lastly, we computed a unit weighted overall biomarker score for each chimpanzee. For this measure, the NPQ values for each marker were converted to standardized *z*-scores, then averaged across all markers to derive a unit weighted average biomarker score (UWA_BM).

## Results

### Patterns of Association Between Age and Biomarkers

We initially tested linear and quadratic associations between age and each individual biomarker and ratio measure using stepwise regression analysis. In the analysis, linear age was step 1, quadratic age was step two, and then the change in F value was evaluated to determine significance. The results are shown in Table 1. The patterns of association between the individual biomarkers and age are shown in Figure 2A-C. Significant positive linear correlations were found between age and APOE4, MAPT, pTau181, pTau231, TREM2, and BD-pTau231. Significant negative linear correlations were also found between age and the ratios of Aβ42/Aβ40 and BD-pTau217/Aβ42.Increasing age was associated with higher values for every marker. Significant quadratic associations were found for Aβ38, Aβ40, Aβ42, GFAP, NfL, PSEN1, pTau217, BD-MAPT, BD-pTau181, and BP-pTau217. For each measure, the NPQ values were higher in elderly and younger chimpanzees relative to middle-aged individuals. Lastly, a significant quadratic association was found between age and the UWA_BM measure.

**Figure 2.**
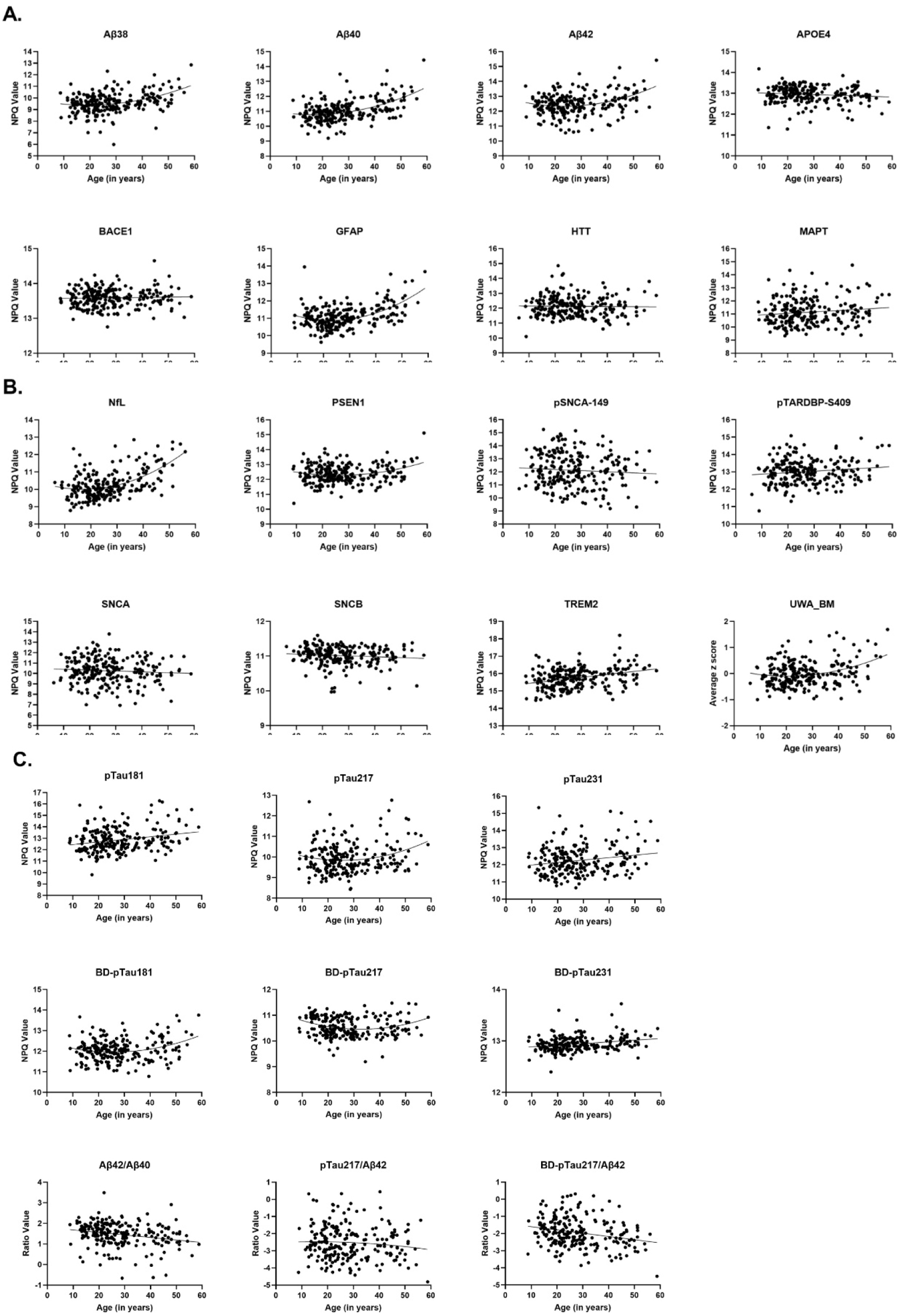
Scatterplots showing the relationships between age and (A.) Aβ38, Aβ40, Aβ42, APOE4, BACE1, GFAP, HTT, MAPT, (B.) NfL, PSEN1, pSNCA-149, pTARDBP-S409, SNCA, SNCB, TREM2, UWA_BM, and (C.) pTau181, pTau217, pTau231, BD-pTau181, BD-pTau217, BD-pTau231, Aβ42/Aβ40, pTau217/Aβ42, and BD-pTau217/Aβ42.

**Table 1.**
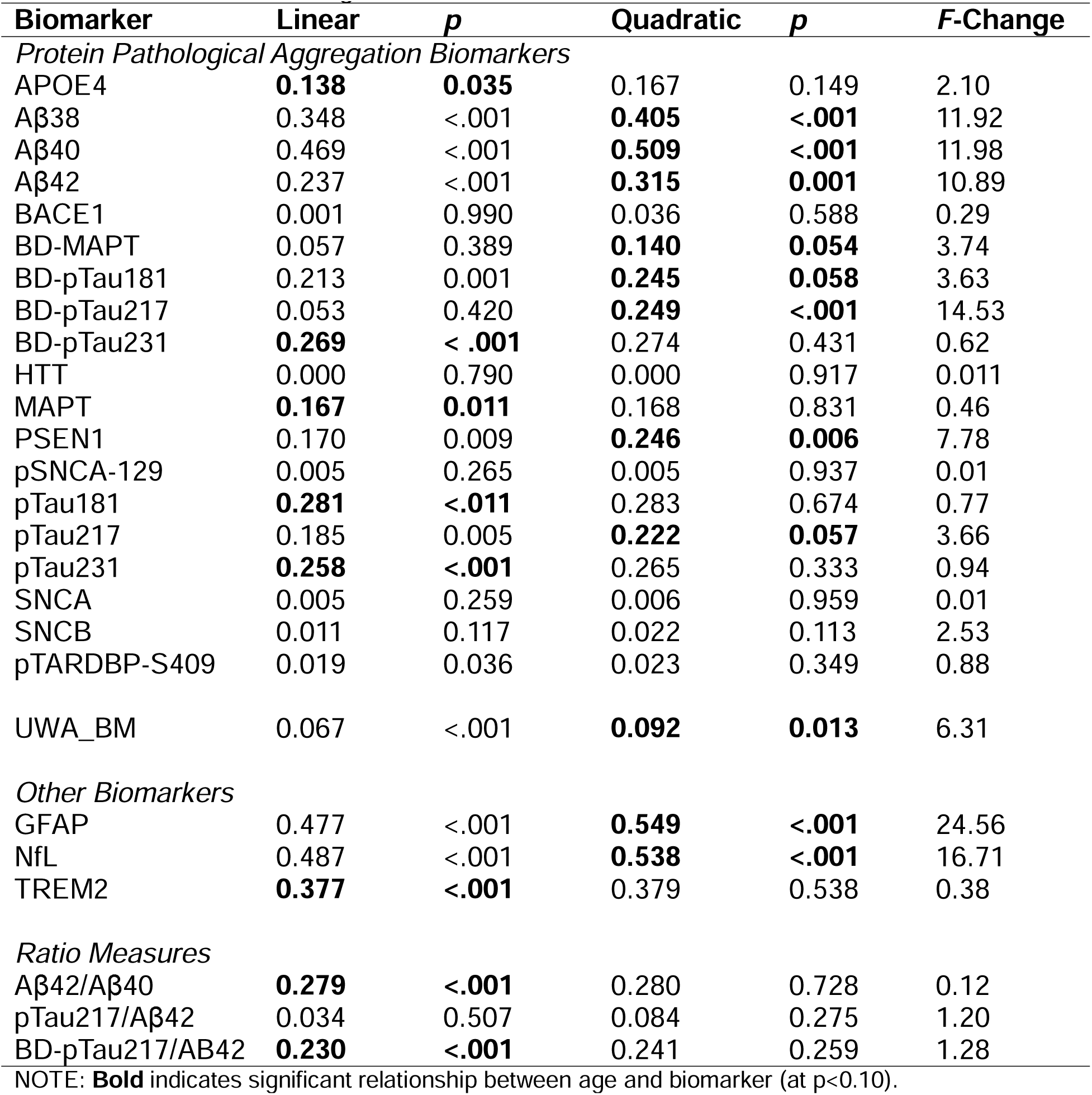
Partial Beta Correlation Coefficients Between Linear and Quadratic Age and.

### Sex and Slope Differences in Age Effects

For these analyses, we performed analysis of covariance (ANCOVA) and built in main effects for sex and age as well as the interaction between sex and age for each outcome measure in the statistical model. Of specific interest were any main effects for sex or the interactions between sex and age. Significant main effects of sex were found for MAPT *F*(1, 229) = 10.99, *p* < .001, pTau181 *F*(1, 230) = 13.236, *p* < .001, pTau217 *F*(1, 229) = 11.437, *p* < .001, pTau231 *F*(1, 229) = 8.024, *p* = .005, pTARDBP-S409 *F*(1, 228)=5.063, *p* = .025 and TREM2 *F*(1, 230)=17.928, *p* < .001. Males had significantly higher NPQ values than females apart from TREM2 (see Figure 3A). Significant two-way interactions between sex and age were found TREM2 *F*(1, 230) = 11.598, *p* < .001, pTau181 *F*(1, 230) = 5.599, *p* = .019, and pTau217 *F*(1, 230) = 4.661, *p* = .033. Shown in Figure 3B are the scatterplots between age and each biomarker for males and females. Females had greater positive slopes in changes as a function of age for pTau181 and pTau217 compared to males. In contrast, males had greater positive slopes in changes as a function of age for TREM2 compared to females.

**Figure 3.**
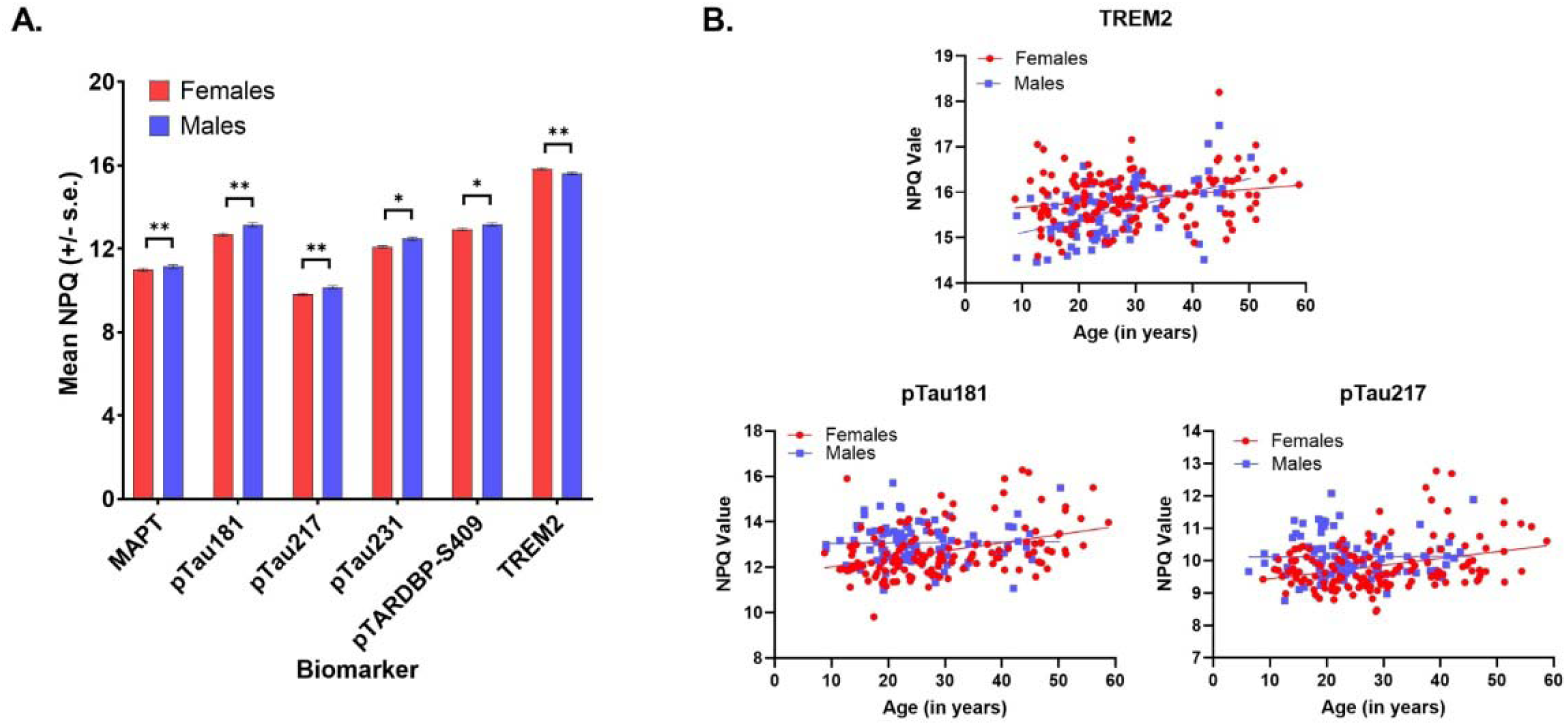
Bar graph (A.) showing significant sex differences in MAPT, pTau181, pTau217, pTau231, pTARDBP-S409, and TREM2 (Mean ± SEM; * indicates p<0.05 and ** indicates p<0.001). Scatterplots (B.) showing the significant sex by age interactions in TREM2, pTau181 and pTau217.

### Longitudinal Analysis

For each biomarker, we used a mixed model analysis of variance with time as a repeated measure and sex as the between-group factor. No significant main effects for sex were found nor were there any interactions between sex and time. Significant main effects for time were found for APOE4 *F*(1,22) =15.08, *p* < .001, Aβ38 *F*(1,22) =4.55, *p* = .045, Aβ40 F(1,22) =14.00, *p* = .001, BD-pTau181 *F*(1,22) = 24.63, *p* < .001, GFAP *F*(1,22) = 6.36, *p* = .019, HTT *F*(1,220) = 8.99, *p* = .007, MAPT *F*(1,22)=13.26, *p* = .001, NfL *F*(1,22)=13.98, *p* = .00, PSEN1 *F*(1,22) = 23.58, *p* < .001, pTau217 *F*(1,22) = 12.45, *p* = .002, pTau231 *F*(1,22) = 8.50, *p* = .008, pTARDBP-S409 *F*(1, 22) = 10.61, *p* = .004 and TREM2 *F*(1,22) = 21.67, *p* < .001. Apart from APOE4, all NPQ values increased with age. The mean NPQ values for time points 1 and 2 for each biomarker are shown in Figure 4. Regarding the ratio measures, a significant main effect for time was found for Aβ42/Aβ40 *F*(1,22) = 9.20, *p* = .006, with higher average Aβ42/Aβ40 at time 1 (M = 1.124, SEM =.010) compared to time 2 (M = 1.079, SEM = .012).

**Figure 4.**
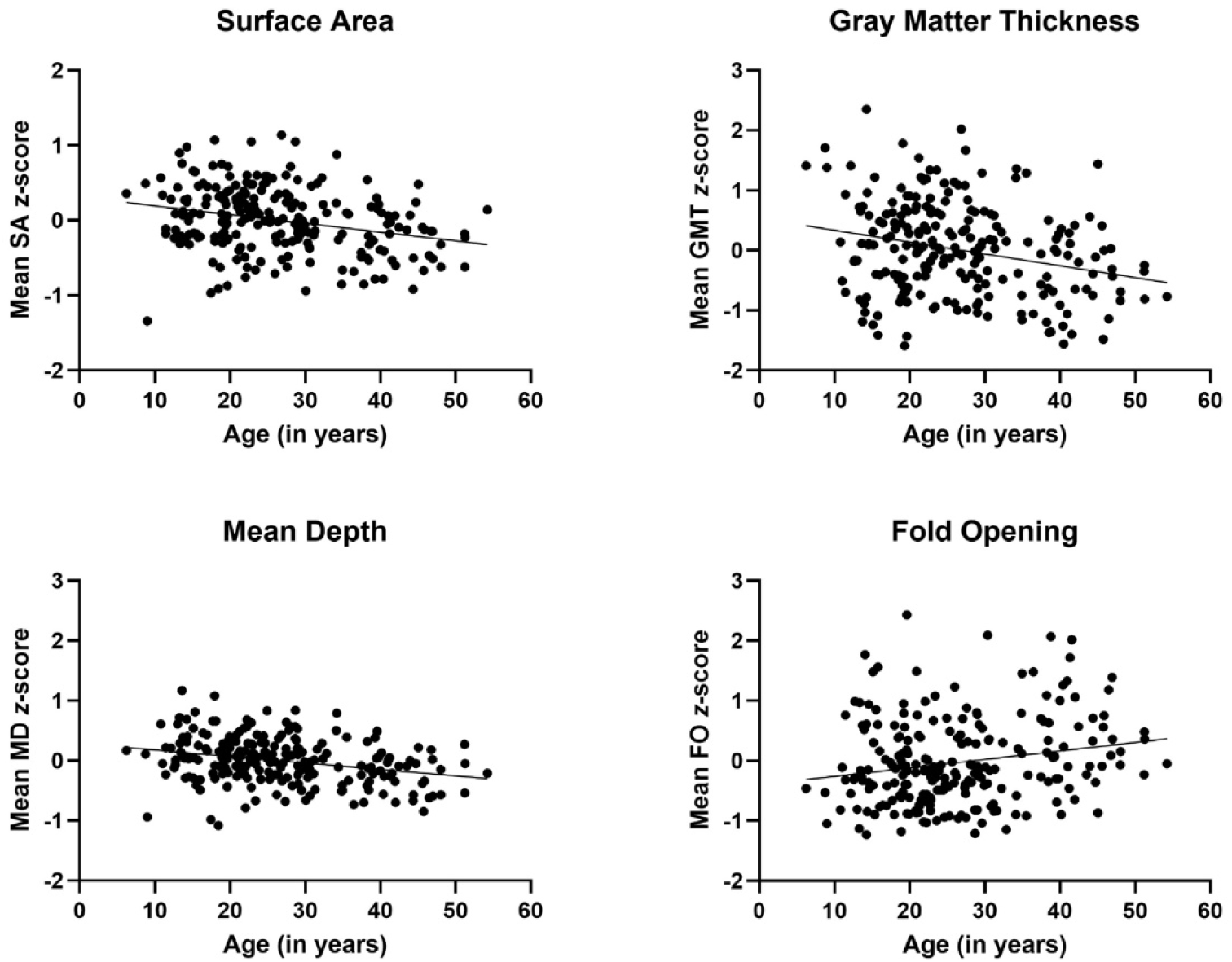
Bar graphs showing the differences in the biomarkers between the two sample time points (Mean ± SEM; * indicates which biomarkers differed)

### Aging and Cortical Atrophy

While controlling for sex, partial correlation coefficients revealed significant negative associations between age and SA (*r* = −.231, *p* < .001), MD (*r* = −.259, *p* < .001), and GMT (*r* = −.267, *p* < .001) (see Figure 5). By contrast, and as predicted, increasing age was associated with higher FO values (*r* = +.213, *p* = .001) (see Figure 5). Thus, as chimpanzee age, their sulci surface areas decline as does their depth. Further, the gray matter surrounding the sulci becomes thinner which, concurrently, results in an expansion of the fold opening or width of the sulci.

**Figure 5.**
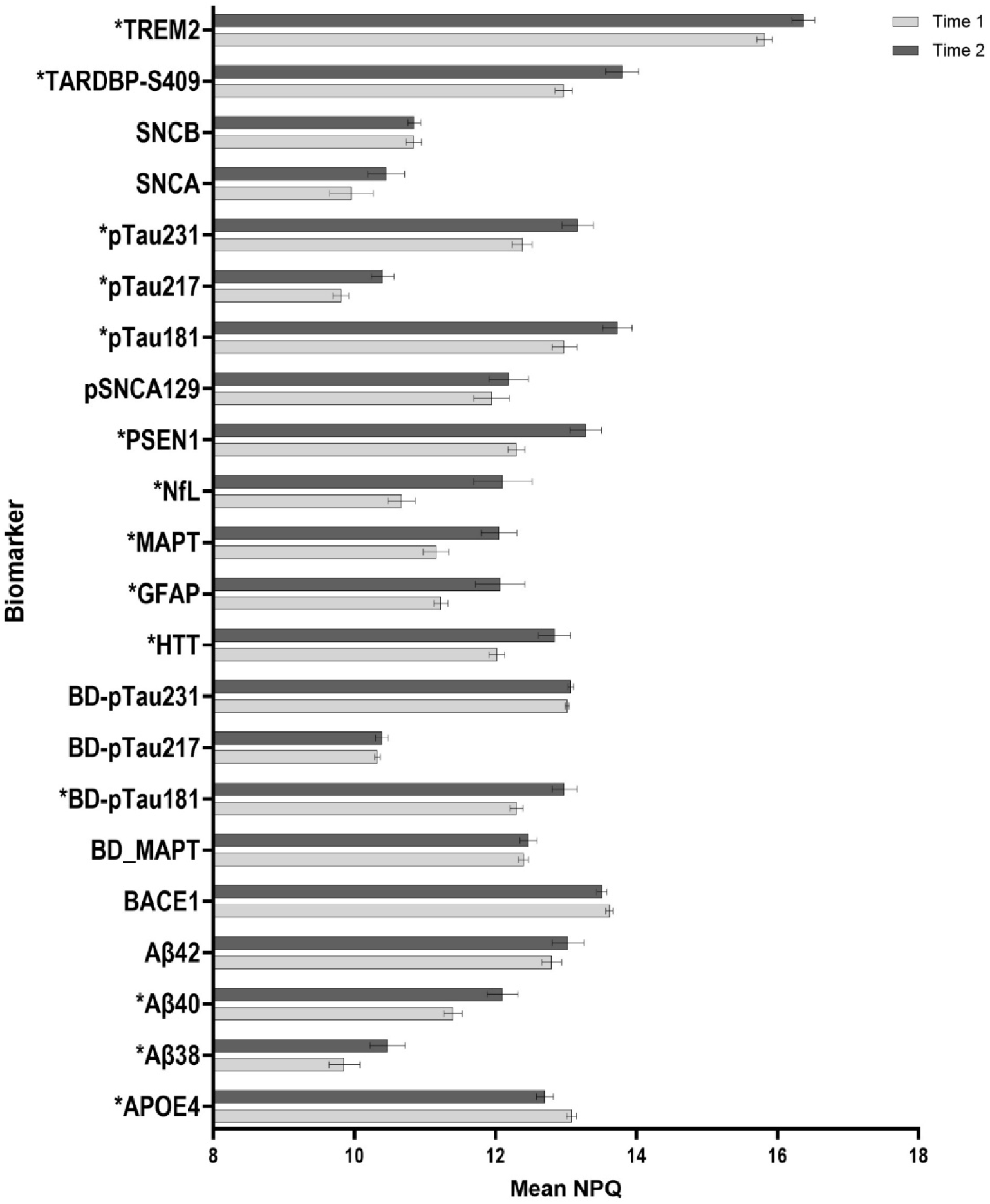
Scatterplots showing significant negative correlations between age and surface area, gray matter thickness, and mean depth), and a significant positive correlation between age and fold opening.

### Patterns of Association Between Biomarkers and Cortical Atrophy

#### Individual Markers

For the individual biomarker measures, the most consistent findings were for Aβ42 and Aβ40 (see Table 2). For these two amyloid biomarkers, increasing values were associated with lower SA, MD and GMT and higher FO values. The remaining significant partial correlations were less consistent across the sulci outcome measures. SA values were inversely correlated with NfL. MD values were inversely associated with Aβ38. GMT was inversely correlated with NfL, while positively correlated with BD-pTau217. Finally, FO was not associated with any additional markers (other than Aβ42 and Aβ40).

**Table 2.** Associations Between Measures of SA, MD, GMT and FO with Each.

| **Biomarker** | **SA** | **MD** | **GMT** | **FO** |
| --- | --- | --- | --- | --- |
| *Protein Pathological Aggregation Biomarkers* |  |  |  |  |
| APOE4 | 0.005 | -0.014 | -0.021 | 0.012 |
| Aβ38 | -0.113 | -0.136* | -0.027 | 0.118 |
| Aβ40 | -0.186** | -0.165 * | -0.164 * | 0.152 * |
| Aβ42 | -0.242 *** | -0.167 * | -0.287 *** | 0.183 ** |
| BACE1 | 0.035 | 0.040 | -0.024 | -0.002 |
| BD-MAPT | 0.068 | -0.013 | 0.023 | 0.059 |
| BD-pTau181 | -0.004 | -0.047 | -0.016 | 0.015 |
| BD-pTau217 | 0.089 | 0.062 | 0.154 * | -0.005 |
| BD-pTau231 | 0.004 | 0.053 | 0.004 | 0.032 |
| HTT | 0.105 | 0.068 | 0.083 | -0.115 |
| MAPT | 0.023 | 0.009 | 0.113 | -0.048 |
| PSEN1 | -0.033 | 0.019 | 0.025 | 0.022 |
| pSNCA-129 | -0.030 | -0.006 | 0.047 | 0.021 |
| pTARDBP-S409 | 0.095 | 0.057 | 0.107 | -0.065 |
| pTau181 | -0.002 | -0.048 | 0.100 | -0.030 |
| pTau217 | -0.011 | -0.039 | 0.101 | 0.005 |
| pTau231 | -0.024 | -0.053 | 0.058 | -0.022 |
| SNCA | -0.006 | -0.001 | 0.025 | 0.017 |
| SNCB | -0.047 | -0.069 | -0.018 | 0.080 |
| UWA_BM | -0.023 | -0.045 | 0.035 | 0.053 |
| *Other Biomarkers* |  |  |  |  |
| GFAP | 0.066 | 0.039 | 0.048 | 0.015 |
| NfL | -0.113 * | -0.128 | -0.138 * | 0.107 |
| TREM2 | -0.075 | -0.029 | 0.087 | 0.023 |
| *Ratio Measures* |  |  |  |  |
| Aβ42/Aβ40 | -0.071 | 0.003 | -0.178 ** | 0.078 |
| pTau217/Aβ42 | 0.158 * | -0.048 | 0.356 *** | -0.173 ** |
| BD-pTau217/Aβ42 | 0.246 *** | -0.012 | 0.435 *** | -0.222 *** |

NOTE: \*p < .05, \*\* p < .01, \*\*\* p < .001

#### Ratio Measures

For the ratio measures, significant negative associations were found between Aβ42/Aβ40 and GMT but not SA, MD or FO. By contrast, pTau217/Aβ42 values were positively associated with SA and GMT and negatively with the FO measure. Similarly, the BD_pTau217/Aβ42 ratio was significantly positively correlated with SA and GMT and negatively with the FO measure. Thus, chimpanzees with higher pTau-217/Aβ42 (both brain-derived and combined) had lower fold opening and higher surface area and gray matter thickness values. See Table 2 for correlation coefficients and Figure 6 for scatterplots.

**Figure 6.**
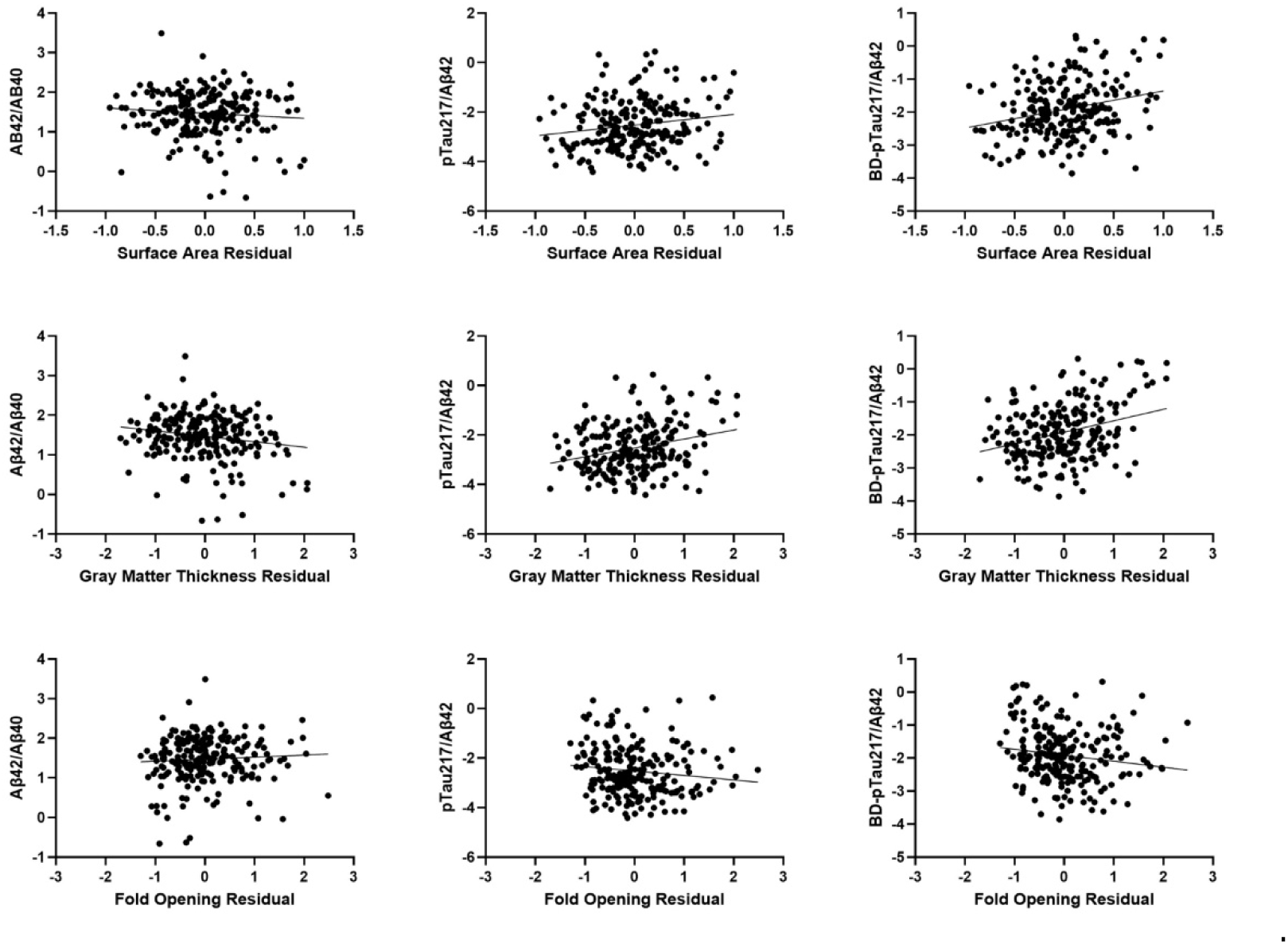
Scatterplots showing the relationship between the biomarker ratio measures and surface area, gray matter thickness, and fold opening residuals.

The associations between the brain measures and pTau217/Aβ42 were confirmed by two follow-up analyses. First, we assessed the correlations between the SA, MD, GMT, and FO values and Aβ42 and pTau217, as well as between Aβ42 and pTau217 while controlling for sex and age. We subsequently tested for differences in the partial *r*-values for Aβ42 and pTau217 for each brain outcome measure using Steiger’s *z*-score. As can be seen in Table 3, significant differences were found between the Aβ42 and pTau217 values for SA, GMT and FO. Further, as reported in humans, Aβ42 showed a negative correlation with the outcome measures while pTau217 showed no association or small positive association, except for FO which is predictably in the opposite direction. The magnitude and difference in the directions of the associations between pTau217 and Aβ42 explains, in part, why their ratio value is significantly associated with the brain phenotypes.

**Table 3.** Summary of Correlation Coefficients Between pTau217 and Aβ42 and Their Differences as Assessed by the Steiger’s z-Score Analysis.

| | pTau217 | Aβ42 | pTau217 + Aβ42 | Steiger's Z-score | p-value |
| --- | --- | --- | --- | --- | --- |
| SA | -0.242 | -0.008 | 0.082 | -2.63 | 0.008 |
| MD | -0.167 | -0.035 | 0.082 | -1.44 | 0.142 |
| GMT | -0.287 | 0.100 | 0.082 | -4.38 | <.001 |
| FO | 0.195 | 0.065 | 0.082 | 2.00 | 0.045 |

In a second analysis, we used stepwise multiple regression to assess the change in proportion of variance accounted for after entering sex, age, the individual pTau217 values, and then the pTau217/Aβ42 ratio values (in that specific order). An *F*-value was computed when entering each variable into the model to assess whether the addition of the variable(s) in the model significantly increased the proportion of variance accounted for (*R*^2^). The full model regression analysis was significant for SA *F*(1, 218) = 15.67, *p* < .001, MD *F*(1, 218) = 10.29, *p* < .001, GMT *F*(1, 218) = 10.16, *p* < .001 and FO *F*(1, 218) = 4.51, *p* = .002. As can be seen in Table 4, sex accounted for a significant proportion of variance for the SA and MD measures. Age accounted for a significant increase in the proportion of variance in all four brain measures. pTau217 alone failed to account for an increase in the proportion of variance for any of the outcome measures; however, the pTau217/Aβ42 ratio accounted for a significant increase in the variance for all four brain outcome measures. We note here that variance inflation factor (VIF) values for the predictor variables for all analyses (see Table 4) were well below acceptable limits (VIF < 3.00) for guarding against collinearity in predictor variables. When considering the brain-derived-pTau217/Aβ42 ratio, as expected, the full model regression analyses were significant for SA *F*(1, 218) = 16.42, *p* < .001, MD *F*(1, 218) = 10.60, *p* < .001, GMT *F*(1, 218) = 10.43, *p* < .001 and FO *F*(1, 218) = 4.50, *p* = .002 (see Table 5). Further, BD-pTau217 alone failed to account for an increase in the proportion of variance for any of the outcome measures; however, the BD-pTau217/Aβ42 ratio accounted for a significant increase in the variance for all brain outcome measures.

**Table 4.** Results of Stepwise Regression Analysis.

| | $R^2$ | $F(1, 218)$ | $p$ | VIF |
| --- | --- | --- | --- | --- |
| *Surface Area* |  |  |  |  |
| Sex | 0.111 | 27.61 | <.001 | 1.13 |
| Age | 0.175 | 17.06 | <.001 | 1.11 |
| pTau217 | 0.175 | 0.01 | 0.910 | 1.99 |
| pTau217/Aβ42 | 0.223 | 13.53 | <.001 | 1.95 |
| *Mean Depth* |  |  |  |  |
| Sex | 0.057 | 13.45 | <.001 | 1.13 |
| Age | 0.134 | 19.56 | <.001 | 1.11 |
| pTau217 | 0.135 | 0.27 | 0.605 | 1.99 |
| pTau217/Aβ42 | 0.159 | 6.06 | 0.015 | 1.95 |
| *Gray Matter Thickness* |  |  |  |  |
| Sex | 0.000 | 0.45 | 0.833 | 1.13 |
| Age | 0.066 | 15.52 | <.001 | 1.11 |
| pTau217 | 0.075 | 2.22 | 0.137 | 1.99 |
| pTau217/Aβ42 | 0.157 | 21.13 | <.001 | 1.95 |
| *Fold Opening* |  |  |  |  |
| Sex | 0.003 | 0.74 | 0.391 | 1.13 |
| Age | 0.044 | 9.42 | 0.002 | 1.11 |
| pTau217 | 0.044 | 0.01 | 0.970 | 1.99 |
| pTau217/Aβ42 | 0.076 | 7.57 | 0.006 | 1.95 |

**Table 5.** Results of Stepwise Regression Analysis (Brain-Derived pTau217).

|  | ***R*<sup>2</sup>** | ***F*(1, 218)** | ***p*** | **VIF** |
| --- | --- | --- | --- | --- |
| *Surface Area* |  |  |  |  |
| Sex | 0.111 | 27.61 | <.001 | 1.13 |
| Age | 0.175 | 17.06 | <.001 | 1.11 |
| BD-pTau217 | 0.185 | 2.69 | 0.103 | 1.99 |
| BD-pTau217 / Aβ42 | 0.223 | 13.19 | <.001 | 1.95 |
| *Mean Depth* |  |  |  |  |
| Sex | 0.057 | 13.45 | <.001 | 1.13 |
| Age | 0.134 | 19.56 | <.001 | 1.11 |
| BD-pTau217 | 0.14 | 1.34 | 0.249 | 1.99 |
| BD-pTau217 / Aβ42 | 0.163 | 6.04 | 0.015 | 1.95 |
| *Gray Matter Thickness* |  |  |  |  |
| Sex | 0.000 | 0.45 | 0.833 | 1.13 |
| Age | 0.066 | 15.52 | <.001 | 1.11 |
| BD-pTau217 | 0.087 | 5.00 | 0.026 | 1.99 |
| BD-pTau217 / Aβ42 | 0.161 | 19.15 | <.001 | 1.95 |
| *Fold Opening* |  |  |  |  |
| Sex | 0.003 | 0.74 | 0.391 | 1.13 |
| Age | 0.044 | 9.42 | 0.002 | 1.11 |
| BD-pTau217 | 0.044 | 0.00 | 0.990 | 1.99 |
| BD-pTau217 / Aβ42 | 0.076 | 7.54 | 0.007 | 1.95 |

## Discussion

Consistent with our hypotheses, the findings from this study indicate that, like in some reports in other nonhuman primates ^8,45–47,62–64^, increasing age is associated with higher levels of amyloid beta, phosphorylated tau, GFAP, NfL, and total tau. We also found a significant association between age and Aβ42/Aβ40 ratio, with younger chimpanzees having higher values, which is consistent with reports in plasma from humans ^65–68^. Given the genetic and biological similarities between humans and chimpanzees, as well as the evidence of AD-related neuropathology in this species, the significant associations between age and the neurodegeneration biomarkers of focus in this study are not all that surprising. That said, the patterns of association between age and biomarkers were more often quadratic than linear with increases in the values becoming incrementally higher beginning at approximately 30 years of age (see scatterplots in Figure 2 and correlation coefficients in Table 1). Huber, et al. ^7^ have suggested that the median lifespan for chimpanzees is approximately 38 years of age which is within the age range of when we found increasing levels of the biomarkers.

The majority of biomarkers associated with age in the cross-sectional analysis also differed in the longitudinal comparisons (see Figure 5). Further, all of the longitudinal changes in the NPQ values between the two testing time points followed the same directional trends as the cross-sectional findings. Thus, the two approaches generally reveal consistent patterns of changes, with older chimpanzees having higher serum biomarker levels (with some exceptions: APOE4 and Aβ42/Aβ40 ratio). Longitudinal studies of aging phenotypes, such as cortical atrophy, cognition, and the measurement of fluid biomarkers are rare or absent in nonhuman primates ^20,22,69^. Indeed, we know of no published data on longitudinal changes in biomarkers of neurodegeneration in nonhuman primates and, therefore, the findings reported here are novel, albeit in a relatively small number of chimpanzees. Follow up studies using additional archived serum samples from across the chimpanzee lifespan may lead to a more precise assessment of the time course of aging-related neurodegenerative changes in this species.

For the individual biomarker measures, broadly speaking, Aβ40 and Aβ42 showed the most consistent associations with the brain measures, while the remaining markers showed less consistent and modest associations with the brain measures. For the sulci measures, higher Aβ40 and Aβ42 values were associated with lower surface areas, mean depth, and gray matter thickness, and higher fold opening values. When considering their ratio, higher values were negatively associated with gray matter thickness. Thus, those chimpanzees with higher Aβ42 relative to Aβ40 had lower gray matter thickness values, a finding that is consistent with at least one previous study in humans measuring cortical thickness ^70^. In humans, the ratio in Aβ42/Aβ40 is a particularly useful clinical indicator of amyloid deposition when quantified using PET imaging or from postmortem analysis of pathology ^71–74^.

Beside the Aβ42/Aβ40 ratio, we also found that increasing pTau217/Aβ42 ratios were associated with higher SA, MD, and GMT values, and lower FO values. These associations were even stronger when using the brain-derived measures of pTau217 when calculating the ratio measure. Thus, chimpanzees with higher pTau217/Aβ42 had thinner gray matter and larger spans between the two sides of the sulci, a finding similar to previous reports in healthy human populations ^75^. Like Aβ42/Aβ40, the pTau217/Aβ42 ratio has been increasingly recognized as an important biomarker that distinguishes individuals with and without Alzheimer’s disease ^76,77^. Additional studies have shown that pTau217/Aβ42 measurements in clinically healthy individuals or patients with mild cognitive impairment (MCI) can be used to predict those that are at risk of developing AD or related dementias ^35,36,78^. In the absence of neuropathological data in relation to either of these measures in chimpanzees, it is difficult to make any firm or definitive conclusions. However, based on these findings, we would postulate that chimpanzees with the lowest Aβ42/Aβ40 and highest pTau-217/Aβ42 values *may* harbor some amyloid plaque or neurofibrillary tangle pathology that, in turn, may impact individual variation in cortical morphology. We would further point out that a significant inverse correlation is found between the pTau-217/Aβ42 and Aβ42/Aβ40 ratio measures (*r* = −.588, *p* < .001) suggesting that these two measures independently capture increasing atrophy in the aging chimpanzee brain, particularly when considering variation in gray matter. Collectively, we believe this is the first evidence demonstrating an association between human clinically validated biomarkers of neurodegeneration, including AD, with brain aging phenotypes in nonhuman primates.

There are some limitations in this study. Notably, the primary analyses were based on cross-sectional serum and MRI data. Though we were careful to utilize serum samples that were collected at the same time as the MRI scans, this did not eliminate the potential for cohort effects. To some extent, the longitudinal findings supported the cross-sectional findings which may suggest the absence of cohort effects, but we still cannot rule them out. Moreover, the longitudinal analysis was based on a limited number of subjects and included samples that were not collected at systematic intervals. A future higher powered, more controlled longitudinal analysis of the biomarkers and MRI data may provide a more accurate representation of age-related changes in chimpanzees. Additionally, we examined phenotypic associations between the biomarkers and SA, MD, GMT and FO measures that were derived by averaging the individual measures across the 23 sulci. This was done to reduce the number of analyses performed and lower the probability of Type I error. However, the atrophy reported here may not be uniform across the entire cortex. Thus, additional studies may want to consider individual sulci or classes of sulci within different lobes or specified brain regions. Third, for the brain-derived pTau17 and pTau231 measures, many of the chimpanzee individual NPQ values were detectable but considered below threshold by the human standard (see Supplemental Table 1). This observation, in and of itself, is interesting because it may imply a fundamental difference between humans and chimpanzees with respect to the amount of tau expressed in the brain, at least as manifest in circulating serum ^79^. It is possible that detection rates would be higher using plasma, and more likely CSF, in the chimpanzees but this will require additional analyses. Finally, in humans, fluid Aβ42/Aβ40 ratio measures decrease as amyloid aggregates in the brain increase ^66,80^. Studies examining both fluid biomarker levels and amyloid pathology in the chimpanzee brain are needed to determine if they too show this inverse relationship.

In summary, the findings reported here are novel in two ways. First, these are the first report of age-related differences in biomarkers of neurodegeneration in chimpanzees. Because of their genetic similarity to humans, these are important reference data for studies on the biology of aging in primates. Second, though there is one previous study that reported significant associations between CSF biomarkers of neurodegeneration and brain morphology in vervet monkeys, we believe the results reported are the first demonstrating associations between measures of cortical atrophy and clinically relevant human biomarkers AD in a nonhuman primate. The combined results suggest that chimpanzees, and perhaps other NHP species, experience age-related changes in the brain which may, in turn, impact aspects of motor or cognitive functions that resemble prodromal or perhaps more advanced stages of neurodegenerative diseases, including AD.

## Funding Sources

This work was supported in part by NIH grants AG067419 (WDH), AG087945 (WDH, EM), AG078411 (MMM), AG087914 (MMM) as well as a grant from Cattlemen for Cancer Research (WDH and MMM) and two French ANR grants: PEPR StratifyAging (ANR-22-PESN-0010) and PEPR PRODROM-ND (ANR-25-APMN-0001). Data were generated in part through the use of the FCCIF-ORION, which received partial support from the National Cancer Institute under grant P30CA016672 to MD Anderson Cancer Center. The research reported here was not directly funded through the P30CA016672 grant to MD Anderson Cancer Center and is not within the scope of such grant.

## Notes

### Competing Interest Statement

The authors have declared no competing interest.

