## Supplemental Table 1 for "Clinically Relevant Biomarkers of Alzheimer’s Disease Are Associated with Age and Cortical Atrophy in Chimpanzees (*Pan troglodytes*)"

| **Supplementary Table 1.** NULISAseq CNS Disease Panel 120 detection rates, as well as each biomarker's linear and quadratic associations with age. | | | | | | |
| --- | --- | --- | --- | --- | --- | --- |
| **Biomarker** | **Threshold** | **Percentage Detection** | **Number of Zero Values or Outliers** | **Association with Age** | | |
|  |  |  |  | **Linear** | **Quadratic** | **N** |
| ACHE | Above Minimum Human Threhold | 100 | 0 | **0.036** | **0.036** | 234 |
| AGRN | Above Minimum Human Threhold | 100 | 3 | **0.172** | **0.186** | 231 |
| ANXA5 | Above Minimum Human Threhold | 100 | 2 | 0.011 | 0.012 | 232 |
| APOE | Above Minimum Human Threhold | High Abundance | 1 | 0.023 | 0.026 | 233 |
| APOE4 | Above Minimum Human Threhold | 100 | 2 | **0.019** | **0.028** | 232 |
| ARSA | Above Minimum Human Threhold | 100 | 0 | 0.004 | 0.005 | 234 |
| Aβ38 | Above Minimum Human Threhold | 98.8 | 2 | **0.121** | **0.164** | 232 |
| Aβ40 | Above Minimum Human Threhold | 98.8 | 2 | **0.22** | **0.259** | 232 |
| Aβ42 | Above Minimum Human Threhold | 96.5 | 2 | **0.056** | **0.099** | 232 |
| BACE1 | Above Minimum Human Threhold | 100 | 0 | 0.001 | 0.001 | 234 |
| BASP1 | Above Minimum Human Threhold | 96.5 | 0 | 0.004 | 0.006 | 234 |
| BD-MAPT | Below Threshold | 1.2 | 7 | 0.003 | 0.02 | 227 |
| BD-pTau-181 | Above Minimum Human Threhold | 100 | 2 | **0.045** | **0.06** | 232 |
| BD-pTau-217 | Below Threshold | 23.3 | 0 | 0.003 | **0.062** | 234 |
| BD-pTau-231 | Below Threshold | 5.8 | 4 | **0.074** | **0.076** | 230 |
| BDNF | Above Minimum Human Threhold | 100 | 1 | 0 | 0.016 | 233 |
| CALB2 | Above Minimum Human Threhold | 100 | 4 | **0.098** | **0.132** | 230 |
| CCL11 | Above Minimum Human Threhold | 100 | 1 | **0.119** | **0.121** | 233 |
| CCL13 | Above Minimum Human Threhold | 100 | 0 | 0 | 0.002 | 234 |
| CCL17 | Above Minimum Human Threhold | 100 | 0 | 0 | 0.012 | 234 |
| CCL2 | Above Minimum Human Threhold | 100 | 2 | 0.007 | 0.01 | 232 |
| CCL22 | Above Minimum Human Threhold | 100 | 0 | 0 | 0.003 | 234 |
| CCL26 | Above Minimum Human Threhold | 100 | 1 | **0.028** | **0.029** | 233 |
| CCL3 | Above Minimum Human Threhold | 100 | 1 | **0.029** | **0.032** | 233 |
| CCL4 | Above Minimum Human Threhold | 100 | 0 | 0.001 | 0.004 | 234 |
| CD40LG | Above Minimum Human Threhold | 100 | 1 | 0.003 | 0.023 | 233 |
| CD63 | Above Minimum Human Threhold | 100 | 0 | **0.066** | **0.066** | 234 |
| CHI3L1 | Above Minimum Human Threhold | 100 | 2 | **0.054** | **0.073** | 232 |
| CHIT1 | Above Minimum Human Threhold | 100 | 0 | **0.246** | **0.286** | 234 |
| CNTN2 | Above Minimum Human Threhold | 98.8 | 1 | **0.04** | **0.042** | 233 |
| CRH | Below Threshold | 17.4 | 2 | 0 | 0.002 | 232 |
| CRP | Above Minimum Human Threhold | High Abundance | 1 | **0.057** | **0.058** | 233 |
| CSF2 | Above Minimum Human Threhold | 100 | 1 | **0.106** | **0.107** | 233 |
| CST3 | Above Minimum Human Threhold | 100 | 0 | **0.161** | **0.208** | 234 |
| CX3CL1 | Above Minimum Human Threhold | 100 | 3 | 0.004 | **0.026** | 231 |
| CXCL1 | Above Minimum Human Threhold | 100 | 0 | **0.025** | **0.027** | 234 |
| CXCL10 | Above Minimum Human Threhold | 100 | 1 | **0.035** | **0.038** | 233 |
| CXCL8 | Above Minimum Human Threhold | 100 | 1 | 0.012 | 0.012 | 233 |
| DDC | Above Minimum Human Threhold | 100 | 0 | 0.004 | **0.067** | 234 |
| ENO2 | Above Minimum Human Threhold | 100 | 1 | **0.054** | **0.055** | 233 |
| FABP3 | Above Minimum Human Threhold | 100 | 0 | **0.033** | **0.065** | 234 |
| FCN2 | Above Minimum Human Threhold | 100 | 1 | **0.024** | **0.026** | 233 |
| FGF2 | Above Minimum Human Threhold | 100 | 0 | 0 | 0 | 234 |
| FLT1 | Above Minimum Human Threhold | 100 | 0 | 0.001 | 0.016 | 234 |
| FOLR1 | Above Minimum Human Threhold | 100 | 0 | **0.04** | **0.041** | 234 |
| GDF15 | Above Minimum Human Threhold | 100 | 0 | **0.347** | **0.347** | 234 |
| GDI1 | Above Minimum Human Threhold | 88.4 | 3 | 0 | 0.014 | 231 |
| GDNF | Above Minimum Human Threhold | 97.7 | 2 | 0 | 0 | 232 |
| GFAP | Above Minimum Human Threhold | 100 | 1 | **0.227** | **0.302** | 232 |
| GOT1 | Above Minimum Human Threhold | 100 | 1 | **0.031** | **0.04** | 232 |
| HBA1 | Above Minimum Human Threhold | 98.8 | 8 | **0.016** | **0.03** | 226 |
| HTT | Above Minimum Human Threhold | 100 | 1 | 0 | 0 | 234 |
| ICAM1 | Not Detectable | 18.6 | 198 |  |  |  |
| IFNG | Above Minimum Human Threhold | 100 | 3 | 0 | 0.002 | 231 |
| IGF1R | Above Minimum Human Threhold | 100 | 1 | **0.067** | **0.074** | 233 |
| IGFBP7 | Above Minimum Human Threhold | 100 | 3 | **0.172** | **0.176** | 231 |
| IL10 | Above Minimum Human Threhold | 100 | 4 | 0.001 | 0.011 | 230 |
| IL12p70 | Above Minimum Human Threhold | 75.6 | 2 | 0.014 | **0.031** | 232 |
| IL13 | Above Minimum Human Threhold | 100 | 1 | 0.002 | 0.002 | 233 |
| IL15 | Above Minimum Human Threhold | 100 | 1 | **0.026** | **0.027** | 233 |
| IL16 | Above Minimum Human Threhold | 100 | 1 | **0.043** | **0.045** | 233 |
| IL17A | Above Minimum Human Threhold | 100 | 3 | 0.001 | 0.004 | 231 |
| IL18 | Above Minimum Human Threhold | 100 | 0 | **0.07** | **0.073** | 234 |
| IL1B | Above Minimum Human Threhold | 76.7 | 1 | 0.014 | 0.015 | 233 |
| IL2 | Above Minimum Human Threhold | 100 | 1 | 0.001 | 0.025 | 233 |
| IL33 | Above Minimum Human Threhold | 100 | 4 | **0.022** | **0.046** | 230 |
| IL4 | Above Minimum Human Threhold | 100 | 0 | 0.015 | 0.018 | 234 |
| IL5 | Above Minimum Human Threhold | 100 | 0 | **0.017** | 0.02 | 234 |
| IL6 | Above Minimum Human Threhold | 100 | 3 | 0 | 0.005 | 231 |
| IL6R | Above Minimum Human Threhold | 100 | 0 | **0.024** | **0.026** | 234 |
| IL7 | Above Minimum Human Threhold | 100 | 0 | 0.005 | 0.007 | 234 |
| IL9 | Above Minimum Human Threhold | 100 | 2 | **0.031** | **0.031** | 232 |
| KDR | Above Minimum Human Threhold | 100 | 0 | 0.013 | 0.024 | 234 |
| KLK6 | Above Minimum Human Threhold | 100 | 3 | **0.184** | **0.196** | 231 |
| MAPT | Above Minimum Human Threhold | 100 | 1 | **0.028** | **0.028** | 233 |
| MDH1 | Below Threshold | 7 | 39 | **0.015** | **0.038** | 195 |
| MME | Above Minimum Human Threhold | 100 | 0 | 0.001 | **0.043** | 234 |
| MSLN | Above Minimum Human Threhold | 100 | 0 | 0.001 | 0.019 | 234 |
| NEFH | Above Minimum Human Threhold | 91.9 | 0 | 0.002 | 0.015 | 234 |
| NEFL | Above Minimum Human Threhold | 100 | 7 | **0.237** | **0.29** | 227 |
| NGF | Above Minimum Human Threhold | 100 | 1 | **0.021** | 0.023 | 233 |
| NPTX1 | Above Minimum Human Threhold | 87.2 | 0 | **0.097** | **0.107** | 234 |
| NPTX2 | Above Minimum Human Threhold | 100 | 0 | **0.037** | **0.051** | 234 |
| NPTXR | Above Minimum Human Threhold | 100 | 0 | 0.015 | 0.018 | 234 |
| NPY | Above Minimum Human Threhold | 100 | 2 | **0.104** | **0.116** | 232 |
| NRGN | Above Minimum Human Threhold | 100 | 1 | **0.021** | **0.026** | 233 |
| Oligo-SNCA | Above Minimum Human Threhold | 27.9 | 1 | 0.011 | 0.011 | 233 |
| PARK7 | Above Minimum Human Threhold | 31.4 | 2 | 0.004 | 0.009 | 232 |
| PDGFRB | Above Minimum Human Threhold | 58.1 | 0 | **0.041** | **0.043** | 234 |
| PDLIM5 | Above Minimum Human Threhold | 100 | 1 | 0.002 | 0.003 | 233 |
| PGF | Above Minimum Human Threhold | 100 | 0 | 0.006 | 0.006 | 234 |
| PGK1 | Above Minimum Human Threhold | 100 | 0 | **0.001** | 0.002 | 234 |
| POSTN | Above Minimum Human Threhold | 100 | 4 | **0.02** | **0.043** | 230 |
| PRDX6 | Above Minimum Human Threhold | 100 | 0 | **0.054** | **0.055** | 234 |
| PSEN1 | Above Minimum Human Threhold | 100 | 1 | **0.029** | **0.061** | 233 |
| pSNCA-129 | Above Minimum Human Threhold | 98.8 | 0 | 0.005 | 0.005 | 234 |
| pTau-181 | Above Minimum Human Threhold | 100 | 0 | **0.079** | **0.08** | 234 |
| pTau-217 | Above Minimum Human Threhold | 97.7 | 0 | **0.034** | **0.049** | 234 |
| pTau-231 | Above Minimum Human Threhold | 100 | 1 | **0.067** | **0.07** | 233 |
| pTDP43-409 | Above Minimum Human Threhold | 64 | 4 | 0.001 | **0.031** | 230 |
| PTN | Below Threshold | 36 | 1 | 0 | 0 | 198 |
| REST | Above Minimum Human Threhold | 100 | 1 | **0.026** | **0.037** | 233 |
| RUVBL2 | Above Minimum Human Threhold | 100 | 0 | **0.039** | **0.042** | 234 |
| S100A12 | Below Threshold | 5.8 | 9 | 0.001 | 0.008 | 225 |
| S100B | Above Minimum Human Threhold | 100 | 0 | 0.001 | 0.003 | 234 |
| SAA1 | Not Detectable | 39.5 | 140 |  |  |  |
| SFRP1 | Above Minimum Human Threhold | 100 | 0 | 0.005 | 0.008 | 234 |
| SFTPD | Above Minimum Human Threhold | 100 | 0 | **0.024** | 0.025 | 234 |
| SLIT2 | Above Minimum Human Threhold | 100 | 0 | 0.003 | **0.037** | 234 |
| SMOC1 | Below Threshold | 7 | 4 | 0.014 | 0.016 | 230 |
| SNAP25 | Below Threshold | 2.3 | 1 | 0.001 | **0.039** | 233 |
| SNCA | Above Minimum Human Threhold | 100 | 0 | 0.005 | 0.006 | 234 |
| SNCB | Above Minimum Human Threhold | 100 | 6 | 0.011 | **0.022** | 228 |
| SOD1 | Above Minimum Human Threhold | 100 | 0 | 0.001 | 0.001 | 234 |
| SQSTM1 | Above Minimum Human Threhold | 100 | 3 | **0.04** | **0.063** | 231 |
| TAFA5 | Above Minimum Human Threhold | 100 | 6 | **0.028** | 0.045 | 228 |
| TARDBP | Above Minimum Human Threhold | 100 | 2 | **0.019** | 0.023 | 232 |
| TEK | Above Minimum Human Threhold | 100 | 0 | **0.079** | **0.09** | 234 |
| TIMP3 | Above Minimum Human Threhold | 100 | 0 | **0.029** | **0.034** | 234 |
| TNF | Above Minimum Human Threhold | 100 | 2 | **0.043** | **0.049** | 232 |
| TREM1 | Above Minimum Human Threhold | 59.3 | 13 | **0.04** | **0.053** | 221 |
| TREM2 | Above Minimum Human Threhold | 100 | 0 | **0.142** | **0.143** | 234 |
| UBB | Above Minimum Human Threhold | 66.3 | 2 | 0.004 | 0.004 | 232 |
| UCHL1 | Below Threshold | 8.1 | 8 | 0 | 0 | 226 |
| VCAM1 | Above Minimum Human Threhold | 100 | 1 | 0.016 | 0.019 | 234 |
| VEGFA | Above Minimum Human Threhold | 100 | 3 | **0.07** | **0.074** | 234 |
| VEGFD | Above Minimum Human Threhold | 100 | 5 | **0.18** | **0.185** | 229 |
| VGF | Above Minimum Human Threhold | 97.7 | 1 | 0.012 | 0.02 | 233 |
| VSNL1 | Above Minimum Human Threhold | 100 | 2 | 0.003 | 0.004 | 232 |
| YWHAG | Below Threshold | 4.7 | 5 | 0 | 0.006 | 229 |
| YWHAZ | Below Threshold | 0 | 9 | 0 | 0.001 | 225 |
| NOTE: **Bold** indicates a significant relationship (p<0.05). | |  |  |  |  |  |
